# TET2 regulates the neuroinflammatory response in microglia

**DOI:** 10.1101/592055

**Authors:** Alejandro Carrillo-Jimenez, Özgen Deniz, Maria Victoria Niklison-Chirou, Rocio Ruiz, Karina Bezerra-Salomão, Vassilis Stratoulias, Rachel Amouroux, Ping Kei Yip, Anna Vilalta, Mathilde Cheray, Alexander Michael Scott-Egerton, Eloy Rivas, Khadija Tayara, Irene García-Domínguez, Juan Garcia-Revilla, Juan Carlos Fernandez-Martin, Ana Maria Espinosa-Oliva, Xianli Shen, Peter StGeorge-Hyslop, Guy Charles Brown, Petra Hajkova, Bertrand Joseph, Jose L. Venero, Miguel Ramos Branco, Miguel Angel Burguillos

## Abstract

Epigenetic mechanisms regulate distinct aspects of the inflammatory response in various immune cell types. Despite the central role for microglia, the resident macrophages of the brain, in neuroinflammation and neurodegeneration little is known about their epigenetic regulation of the inflammatory response. Here, we show that Ten-eleven translocation 2 (TET2) methylcytosine dioxygenase expression is increased in microglia upon stimulation with various inflammogens through a NF-κB-dependent pathway. We found that TET2 regulates early gene transcriptional changes that lead to early metabolic alterations, as well as a later inflammatory response independently of its 5mC oxidation activity at the affected genes. We further show that TET2 regulates the proinflammatory response in microglia induced by intraperitoneal injection of LPS *in vivo*. We observed that microglia associated to amyloid β plaques, recently defined as disease-associated microglia, expressed TET2 in brain tissue from individuals with Alzheimer’s disease (AD) and in 5×FAD mice. Collectively, our findings show that TET2 plays an important role in the microglial inflammatory response, and suggest TET2 as a potential target to combat neurodegenerative brain disorders.

## Introduction

Microglia, the resident immune cells in the central nervous system (CNS), are key players in maintaining homeostasis in the brain. Microglia play a wide variety of roles both under physiological and pathological conditions. In the healthy brain, microglia are responsible for neuronal activity-dependent synapse pruning through engulfing presynaptic inputs early during postnatal development (Schafer et al., 2012) (Wu et al., 2015). Upon neuronal injury or infection, microglia become rapid responders that initiate an innate inflammatory response (Hanisch and Kettenmann, 2007). If the inflammatory response is exaggerated or becomes chronic, it results in a detrimental response for the surrounding neuronal population. This excessive inflammatory response occurs in many neurodegenerative disorders such as Parkinson’s (PD) and Alzheimer’s diseases (AD) (Ransohoff, 2016),(Perry and Holmes, 2014),(Burguillos et al., 2011),(Abeliovich and Gitler, 2016), as well as in neurological conditions such as ischemic stroke (Lambertsen et al., 2012),(Burguillos et al., 2015). However, the mechanisms that trigger this exacerbated response in these different neurodegenerative disorders are still not clear.

In PD or AD, only a minor subset of patients has a genetic mutation responsible for disease development (e.g., *SNCA, PINK1* and *PARK2* mutations in PD (Abeliovich and Gitler, 2016)(Pickrell and Youle, 2015), and *APP, PSEN1, PSEN2* in AD (Van Cauwenberghe et al., 2016)). The majority of cases appear to be a consequence of a combination of genetic predisposition and the exposure of environmental risk factors. The identification of mutations in innate immunity-related genes that confer higher risk of developing neurodegenerative diseases (*TREM2, CD33, CR1*, etc.), supports the idea of microglia playing a key role driving the pathogenesis in these diseases (Malik et al., 2015). Hence, epigenetic mechanisms are prime candidates for mediating environmentally driven alterations to immune homeostasis that can be inherited across cell division. Indeed, the contribution of epigenetic modifications to neurodegenerative diseases such as PD (Wüllner et al., 2016) (Park et al., 2015) and AD (Watson et al., 2016), (Phipps et al., 2016) has been addressed in a number of studies. However, despite the key role of microglia in the neuroinflammatory response in those neurodegenerative diseases, little is known about the epigenetic regulation of the inflammatory response in these cells.

Major epigenetic mechanisms include post-translational modification of histones (e.g., methylation, acetylation), DNA methylation at CpG dinucleotides, and regulation by non-coding RNAs (Bonasio et al., 2010). Interestingly, the age-dependent increase in microglial IL-1β levels is associated with DNA hypomethylation within the *IL-1β* promoter (Matt et al., 2016), which is seemingly driven by the age-dependent loss of microglial SIRT1, a NAD-dependent deacetylase that can regulate the activity of DNA methyltransferase 1 (DNMT1) (Cho et al., 2015). DNA methylation could, therefore, play key roles in regulating the inflammatory state in microglia. Importantly, DNA methylation can be removed by the action of Ten-eleven Translocation (TET) enzymes, which are dioxygenases that catalyze the oxidation of 5-methylcytosine (5mC) into 5-hydroxymethylcytosine (5hmC) and other oxidative derivatives (Branco et al., 2012). Recently, TETs have been shown to play various roles in the physiology of immune cells. For instance, TET2 and TET3 are responsible for the development and proliferation of CD4+CD8+ double-positive thymocytes into invariant natural killer T cells (iNKT cells) (Tsagaratou et al., 2016). In T helper cells, TET2 mediates demethylation of putative regulatory elements in genes associated with T cell differentiation, regulating the Th1 and Th17 cytokine expression *in vitro* (Ichiyama et al., 2015). TET2 also regulates the inflammatory response in dendritic cells and bone marrow-derived macrophages independently of its dioxygenase activity (Zhang et al., 2015). The latter study showed that TET2 is an important player in the resolution of inflammation by repressing IL-6 expression through recruitment of HDAC2, a histone deacetylase, into the *Il-6* promoter. A role for TET2 during the resolution of inflammation has also been described in bone-derived macrophages (Cull et al., 2017).

Here we investigated the role of TET enzymes in the inflammatory response in microglia cells upon Toll-like receptor 4 (TLR-4) activation. We found that TET2 is an NF-κB-responsive gene that regulates both early transcription (after only a few hours) of genes affecting several pathways (including control of the immune response and cell cycle) and the late inflammatory response. We confirmed *in vivo* (in an inflammatory model induced by intraperitoneal injection of lipopolysaccharide -LPS-) that TET2 regulates the proinflammatory response in microglia cells. Furthermore, we show that TET2 is expressed in microglia close to β-amyloid plaques in a 5xFAD neurodegenerative disease model. Finally, we analyze the expression of TET2 in microglia cells in different areas in the brain of three AD patients. All these results highlight the potential of TET2 as a novel drug target for neurodegenerative diseases, including AD.

## Results

### TLR activation in microglia induces upregulation of Tet2 expression

To assess the effect of TLR-4 activation on the expression of TET enzymes in microglia, we treated the murine BV2 microglial cell line with LPS. We found that LPS (1 µg/ml) induced both an early 2h (Figure 1A) and sustained 24h (Figure 1B) upregulation of *Tet2* expression. However, the expression of the other two members of the TET family (TET1 and TET3) either did not vary upon LPS treatment (*Tet3*; Figure 1A) or was not detectable (*Tet1*; not shown) before or after LPS treatment. In concordance with the RNA data, we detected an increase of TET2 expression at the protein level 6h after LPS treatment (Figure 1C). Interestingly, a lower dose of LPS (0.1 μg/ml) was also able to promote *Tet2* expression as early as 2h after treatment (Figure 1D). To rule out the possibility that LPS-induced TET2 expression might be due to the transformed origin of our murine microglia cell line (Butovsky et al., 2014), we analyzed the expression of *Tet2* in primary adult and postnatal primary microglia cells from mouse and rat origin 6h after LPS treatment, and obtained similar results to those seen in BV2 microglia cells (Figure 1E). Notably, we also observed a mild but significant upregulation of *TET2* in human microglia cells (CHME3 human microglia cell line) (Figure 1E). We further validated our observations by analyzing RNA sequencing (RNA-seq) data from postnatal primary microglia cell culture experiments (Janova et al., 2016), which showed that *Tet2* expression was increased both at low and high levels of LPS treatment, as well as after treatment with fibronectin, which also promotes inflammation (Figure S1A). Fibronectin-mediated regulation of *Tet2* suggests that *Tet2* upregulation might not only be driven by TLR-4 activation. For this reason, we challenged our BV2 cells with Lipoteichoic acid (LTA), a known TLR-2 ligand, and observed a similar pattern in the expression of *Tet2* and *Tet3* to that seen in LPS-treated cells (Figure S1B and S1C). These data show that multiple TLR agonists drive *Tet2* upregulation in microglia from different species.

**Figure 1:**
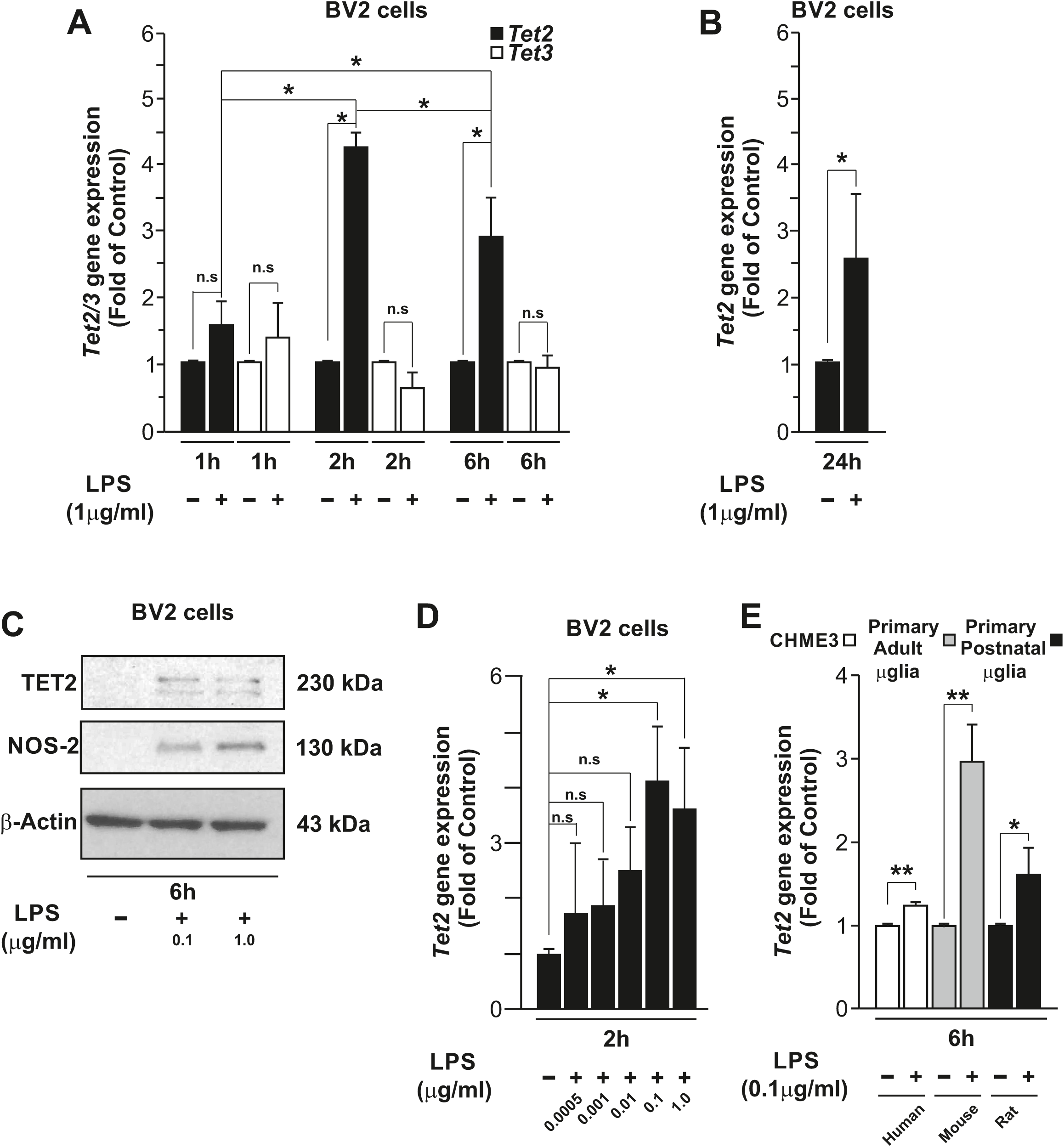
LPS induces an early and sustained expression of *Tet2* in microglia cells. *Tet2* and *Tet3* expression in BV2 microglia cells treated with LPS (1 μg/ml) at 1, 2 and 6h (A) and *Tet2* expression after 24 h treatment with LPS (1 μg/ml) (B). Representative immunoblot showing increase of TET2 and NOS-2 expression (a positive control of microglia activation) at 6h after LPS treatment (0.1μg/ml and 1μg/ml) in BV2 cells (C). *Tet2* LPS-induced dose-response in BV2 microglial cells at 2h (D). *Tet2* gene expression after 6h LPS (0.1μg/ml) treatment in human microglial cell line (CHME3), primary adult microglia in mouse and primary postnatal microglia culture in rat (E). Statistical analysis was performed using one-way analysis of variance (ANOVA) with Scheffe (A, B) and LSD (D) corrections or two-tailed Student’s t-test (E). Data shown are mean ± s.d. of three (A, D and E) and five (B), independent experiments. \**P* < 0.05, \*\**P* < 0.01. See also Figure S1.

### NF-κB p65 mediates LPS-induced Tet2 expression

We then sought to investigate the mechanisms responsible for the transcriptional regulation of *Tet2* upon TLR-4 activation. We first took advantage of published chromatin immunoprecipitation sequencing (ChIP-seq) data on the TLR-4-induced “enhancer landscape” in macrophages (Kaikkonen et al., 2013). We used these data as a model for TLR-4-induced regulatory events that may also be occurring in microglia cells. We mapped data for the activating histone mark H3K27ac, as well as various transcription factors, and visually inspected the promoter and upstream regions of *Tet2* (profiles in Figure 2A and Figure S2A). In untreated macrophages, H3K27ac was enriched both at the promoter region of *Tet2* and a region lying 40kb upstream, thus constituting a putative distal enhancer element (Figure 2A). Notably, the levels of H3K27ac increased in the upstream region (E1 and E2) after 1-hour treatment with KLA (a TLR-4 agonist), and this was concomitant with the recruitment of p65 to both the promoter and upstream regions upon KLA treatment (Figure 2C). In contrast, the binding of CEBPA and PU.1 was largely unaffected by KLA treatment (Figure S2A), suggesting that p65 is a major driver of TLR-4-dependent activation of *Tet2*. To test whether similar patterns can be observed in BV2 cells, we performed ChIP followed by quantitative polymerase chain reaction (qPCR) analysis at the promoter and upstream regions of *Tet2*. In concordance with the results obtained from bone marrow derived macrophages (Figure 2A), BV2 cells are enriched for H3K27ac at the *Tet2* promoter and at the putative regulatory region upstream of *Tet2* (Figure 2B). Interestingly, H3K27ac levels specifically increased in the upstream region upon LPS treatment (Figure 2B). Moreover, we detected enrichment for H3K4me1, a histone mark associated with both poised and active enhancer elements, providing support for the upstream region being an enhancer element that becomes active upon TLR-4 activation (Creyghton et al., 2010). We then analysed p65 enrichment in the promoter and enhancer regions of *Tet2* upon LPS treatment in BV2 cells, and observed a clear increase in p65 binding at the enhancer region, whereas LPS more subtly modulated the binding of p65 at the promoter region (Figure 2D). As positive controls for LPS-dependent p65 binding, we analysed the promoter regions of NF-κB Inhibitor Alpha (*NFκBia*) and *Il-1β* (Figure S2B). These results suggest a potential role of p65 in*Tet2* expression through binding to an upstream enhancer element, increasing its activity, which is reflected by the higher H3K27ac levels in LPS-treated cells.

**Figure 2:**
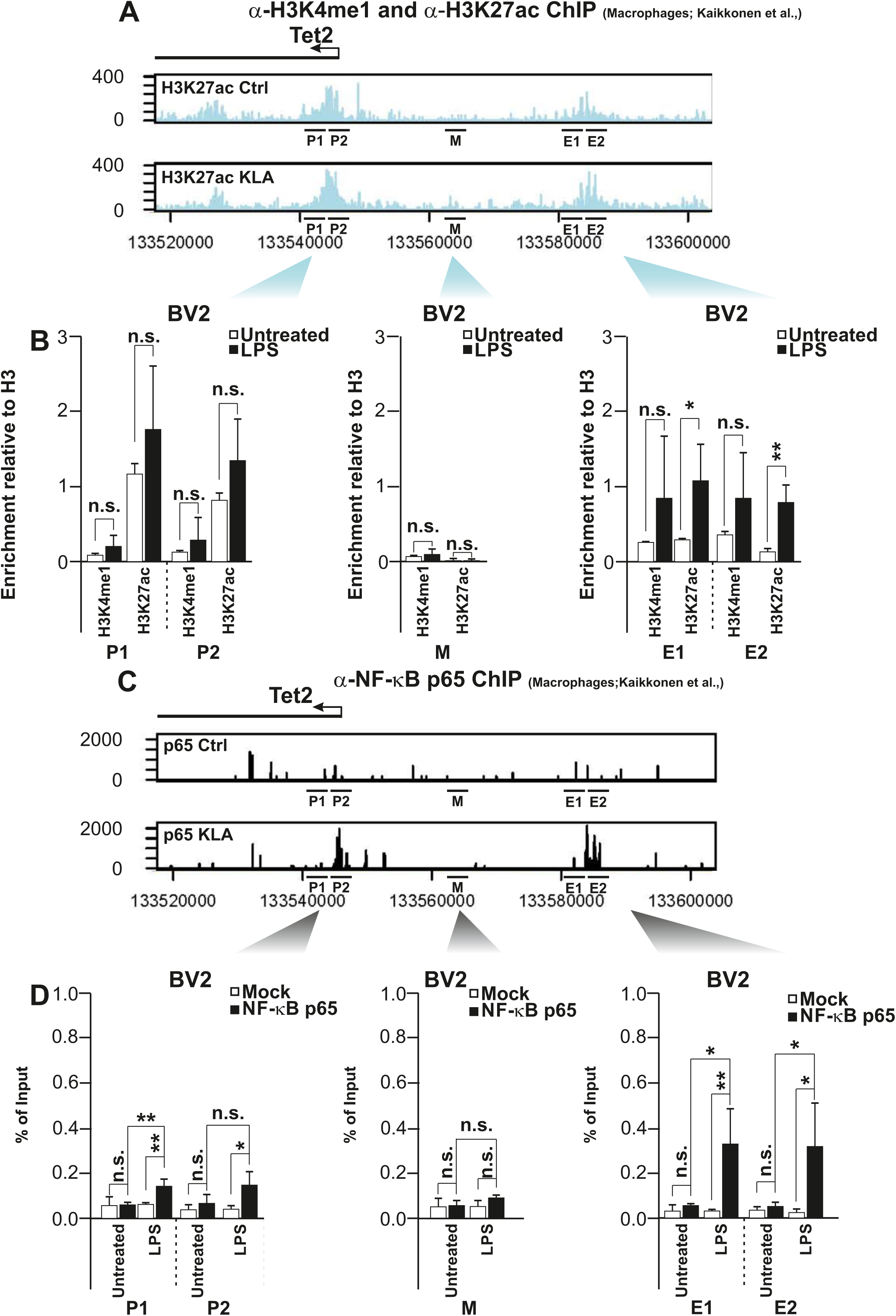
NF-κBp65 regulates LPS-induced *Tet2* transcription. Profile of H3K27ac marking at the *Tet2* promoter and upstream regions after 1h treatment of macrophages with KLA, generated from published ChIP-seq data (Kaikkonen et al. 2013) (A). ChIP-qPCR results for H3K27ac and H3K4me1 at the promoter (P1 and P2), middle (M) and enhancer (E1 and E2) regions in BV2 cells treated with LPS (1μg/ml) (B). Profile of NF-κB p65 binding at the *Tet2* promoter and upstream regions after 1h treatment of macrophages with KLA, generated from published ChIP-seq data (Kaikkonen et al., 2013)(C). ChIP-qPCR results for NF-κB p65 in the same regions as in (B) in BV2 cells treated with LPS (1μg/ml) (D). *Tet2* middle region (M) represents a region between the promoter and the enhancer used as negative control for ChIP. Data represented as mean ± s.d. The results in B are the average of three (for H3K4me1) and four (for H3K27ac) independent experiments. The results in D are the average of 3 independent experiments. Statistical analysis was performed using two-tailed Student’s t-test. See also Figure S2.

To test the functional relevance of increased p65 binding to the upstream region of *Tet2* in regulating *Tet2* expression upon LPS treatment, we pre-treated BV2 cells for 1 hour with wedelolactone, an inhibitor of the IKK complex, followed by treatment with LPS for 6h. We used *Il-1β* as a positive control, as its transcription has been shown to be regulated by NF-κB (Cogswell et al., 1994) (Figure S2C). In line with our ChIP-qPCR data, LPS-induced expression of *Tet2* was abrogated in the presence of wedelolactone (Figure S2D), suggesting that the NF-κB complex plays a role in regulating *Tet2* expression (either directly or indirectly) upon LPS treatment.

### TET2 helps to drive the expression of genes induced by TLR-4 stimulation

Given the reported involvement of TET2 in the regulation of immune functions, we asked whether it also plays a role during the neuroinflammatory response. For this purpose, we depleted *Tet2* in BV2 microglia cells using a specific siRNA against it 48h prior to LPS treatment (Figure 3A). To confirm that TET2 depletion resulted in decreased enzymatic activity, we measured global 5hmC levels by mass-spectrometry, and observed a significant 5hmC reduction in siTET2 cells when compared to a non-targeting control (Figure 3B). Interestingly, LPS treatment did not change global 5hmC levels, despite the increase in TET2 expression. We then performed RNA-seq on TET2-depleted cells, before and after 3h of LPS treatment, and compared them against a non-targeting control. We first confirmed that the expression pattern in our BV2 cells after LPS treatment is very similar to previously published RNA-seq data from LPS-treated primary microglia cells in (Janova et al., 2016) (Figure S3A). To identify genes whose activation/repression during LPS treatment depends on TET2, we intersected LPS-regulated genes with TET2-regulated ones. Out of 1,565 genes that were upregulated by LPS, 140 (9%) had reduced expression in TET2-depleted cells (Figure S3B). Conversely, out of 1,110 genes repressed by LPS, 38 (3%) had increased expression in TET2-depleted cells (Figure S3B). Both groups of TET2-regulated genes displayed an impaired response to LPS in *Tet2* knockdown cells, as judged by significant differences in the expression fold change upon LPS treatment (Figure 3C). Using qPCR analysis, we validated several of the gene expression changes mediated by TET2 depletion in LPS-treated cells (Figure S3C). Gene ontology analyses revealed that the 140 siTET2-downregulated genes are mainly associated with the control of the innate immune response, including the response to interferon-β (also known as Type I-IFN response), whereas the 38 siTET2-upregulated genes are associated with cell cycle regulation (Figure 3D). A manual classification of gene function based on literature searches confirmed an enrichment for inflammatory and cell cycle related genes in TET2-regulated targets, followed by genes related to intracellular signaling and transcription factors (Figure 3E). Interestingly, gene ontology analysis on siTET2-downregulated genes that are not modified by LPS treatment, revealed a similar enrichment for genes involved in the immune response (Figure 3F). This result suggests that the effect of TET2 over the inflammatory response is not unique to LPS treatment but it can also potentially affect other immune signaling pathways.

**Figure 3:**
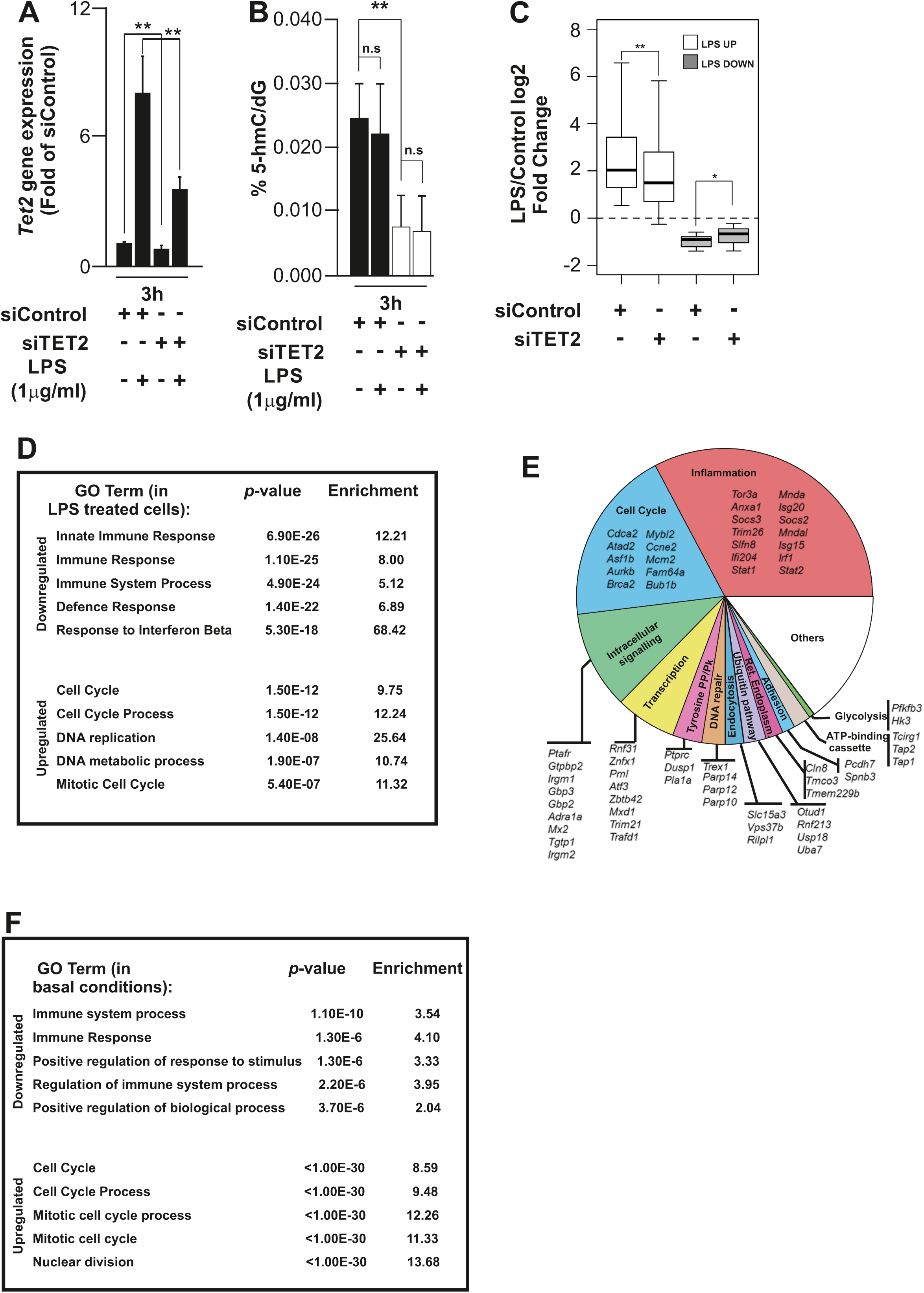
TET2 regulates signaling pathways induced by LPS. Effect of siRNA Tet2 knockdown on *Tet2* gene expression (A) and on the global levels of 5hmC (B) with or without 3h treatment with LPS (1μg/ml). Expression change of LPS-responsive TET2-regulated genes (see Venn diagrams in Figure S3B) upon LPS treatment in siRNA control and TET2 treated cells (C). Table representing gene ontology (GO) analysis of all genes affected by *Tet2* knockdown after LPS treatment (D). Manual annotation representing different functional groups (and some examples of the genes) affected by *Tet2* knockdown after LPS treatment (E). Manual annotation representing different functional groups (and some examples of the genes) affected by *Tet2* knockdown under basal conditions (F). Data shown are represented as mean ± s.d. from five (A), three (B,C) independent experiments. Two-tailed Student’s t-test. \**P* < 0.05, \*\**P* < 0.01. See also Figure S3.

To rule out the possibility that the observed gene deregulation was due to increased cell death, we performed FACS analyses of control and TET2-depleted cell. We could not find any indication of induction of cell death (Figure S3D-G) or major change in morphology (Figure S3H), suggesting a direct effect of TET2 over many of those genes upon treatment.

Altogether, our results suggest that TET2 not only plays a role in the regulation of the inflammatory response, but also in other aspects of microglia physiology, such as cell cycle regulation.

### TET2 does not affect DNA methylation levels at target genes

TET2 has been shown to act via both 5mC oxidation and catalytic-independent mechanisms, such as recruitment of epigenetic modifiers (Ichiyama et al., 2015), (Zhang et al., 2015). We therefore asked whether the regulatory effect of TET2 on LPS-driven gene expression in microglia was dependent on its catalytic activity. We first analyzed global 5mC levels by mass spectrometry in LPS-treated cells, comparing TET2-depleted cells with controls (Figure S4A). Neither LPS treatment nor knockdown of TET2 resulted in a significant change in global 5mC levels (Figure S4A) despite the fact that global 5hmC levels were altered in siTET2 cells (Figure 3B). In order to determine whether TET2-mediated 5mC oxidation occurs in a locus-specific manner, we therefore used oxidative bisulfite sequencing (oxBS-seq) (de la Rica et al., 2016)(Booth et al., 2012) and measured 5mC and 5hmC levels at the promoters of six target genes whose expression levels were altered by TET2 knockdown (Figure 4A-F and Figure S4B-G). No significant changes were detected in the levels of 5mC (Figure 4A-F). In 5hmC levels we observed some statistically significant albeit very minor differences (Figure S4C). These results suggest that the action of TET2 on these genes does not involve its catalytic activity at these gene promoters. In line with this, our mass spectrometry data show that LPS treatment does not induce global changes in 5hmC levels, suggesting that increased TET2 catalytic activity is not necessary to mediate gene expression changes upon LPS treatment.

**Figure 4:**
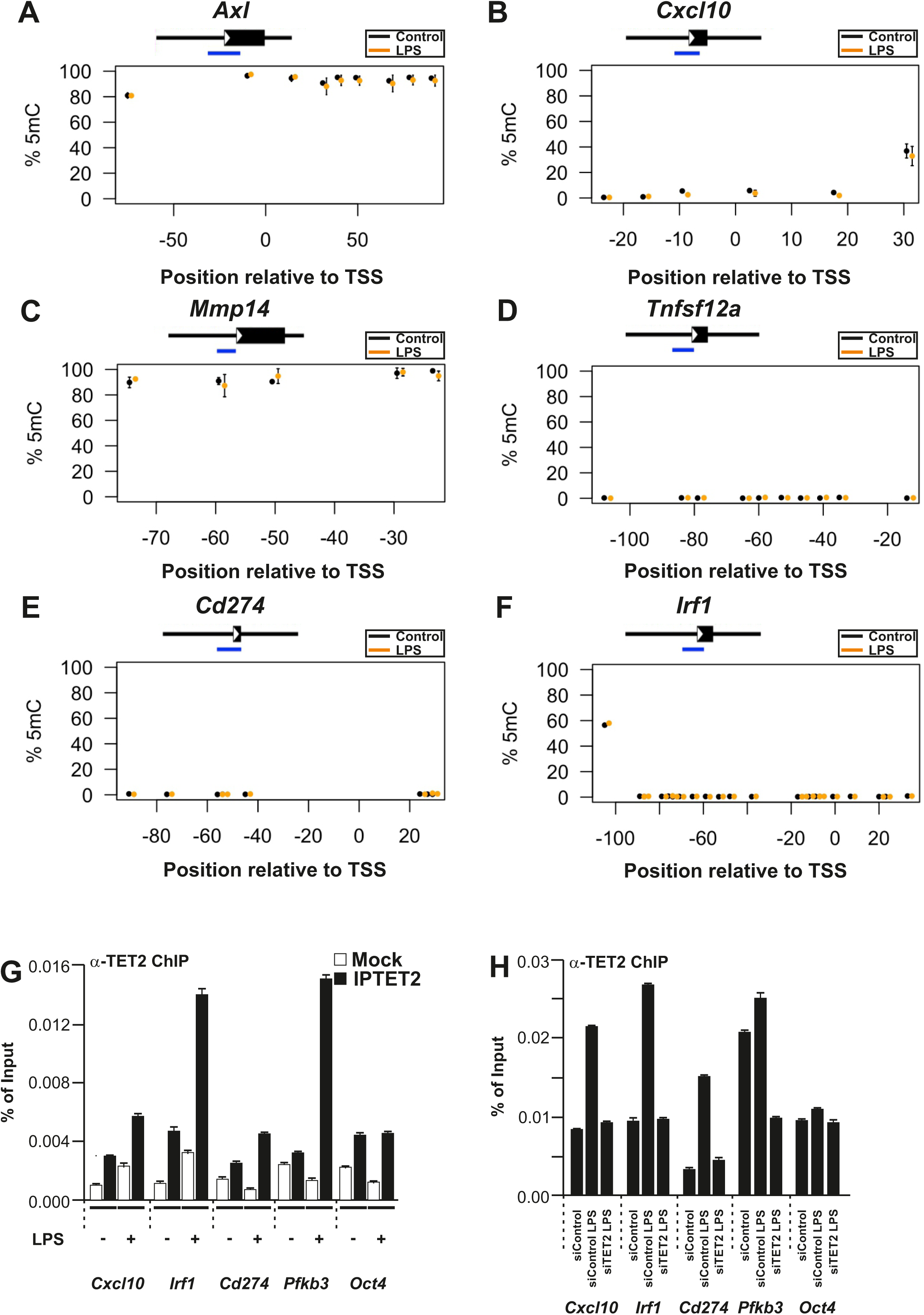
TET2 binds to the promoter regions of several target genes. Quantification of 5mC levels by oxBS-seq of the promoters of target genes after 3h LPS (1μg/ml) treatment (A-F). Blue bars indicate the position of the analysed amplicons. TET2 ChIP for target genes after 3h LPS treatment (G). TET2 ChIP of the genes after 3h treatment with LPS (1μg/ml) in control and TET2-depleted cells (H). *Oct4* was used as a negative control. Data shown in A-H represent the mean ± s.d of two technical replicates. Results from an independent biological replicate of data in H are shown in Figure S4H. Statistical analysis was performed using two-way ANOVA with a Tukey correction tacking into account all CpGs for A-F. See also Figure S4.

The effect of TET2 on the expression of selected genes could also be explained by indirect effects. To test whether TET2 was bound to the promoters of genes affected by TET2 depletion, we performed TET2 ChIP-qPCR. Our data show that, indeed, TET2 binding increases substantially at target genes upon LPS treatment, whereas a control locus (*Oct4*) shows no alterations (Figure 4G). Importantly, LPS-driven TET2 recruitment can be reversed upon knockdown of *Tet2* (Figure 4H and Figure S4H). These results suggest that TET2 acts directly on these genes but that its effect upon LPS is not predominantly mediated through its catalytic activity.

### TET2 regulates the “classical” inflammatory response and the metabolic reprogramming induced by LPS

Our RNA-seq data suggested that TET2 plays a role in the LPS-induced inflammatory response, in particular the response to interferon β (or Type I IFN response) (Figure 3E). It was previously shown that TET2 is required for the repression of IL-6 upon LPS treatment in peripheral macrophages to ensure termination of inflammation (Zhang et al., 2015). IL-6 expression was unchanged in our RNA-seq, which was performed 3h after LPS treatment. Given that 3h is too early for the inflammatory resolution process to start, we analyzed the expression levels of IL-6 and other players related to the “classical” pro-inflammatory response (in this case IL-1β and NOS-2 expression) at later time points upon LPS treatment (Figure 5A-5E). We observed in BV2 cells that, while there was no difference at 3h after LPS treatment, the expression of *Il-1β, Il-6* and *Nos-2* was reduced at 6 and 24h post-LPS treatment upon TET2 depletion (Figure 5A-5C). Knockdown of TET2 led to significant less IL-6 release into the media upon LPS treatment at 6 and 24h in BV2 cells (Figure 5D).

**Figure 5:**
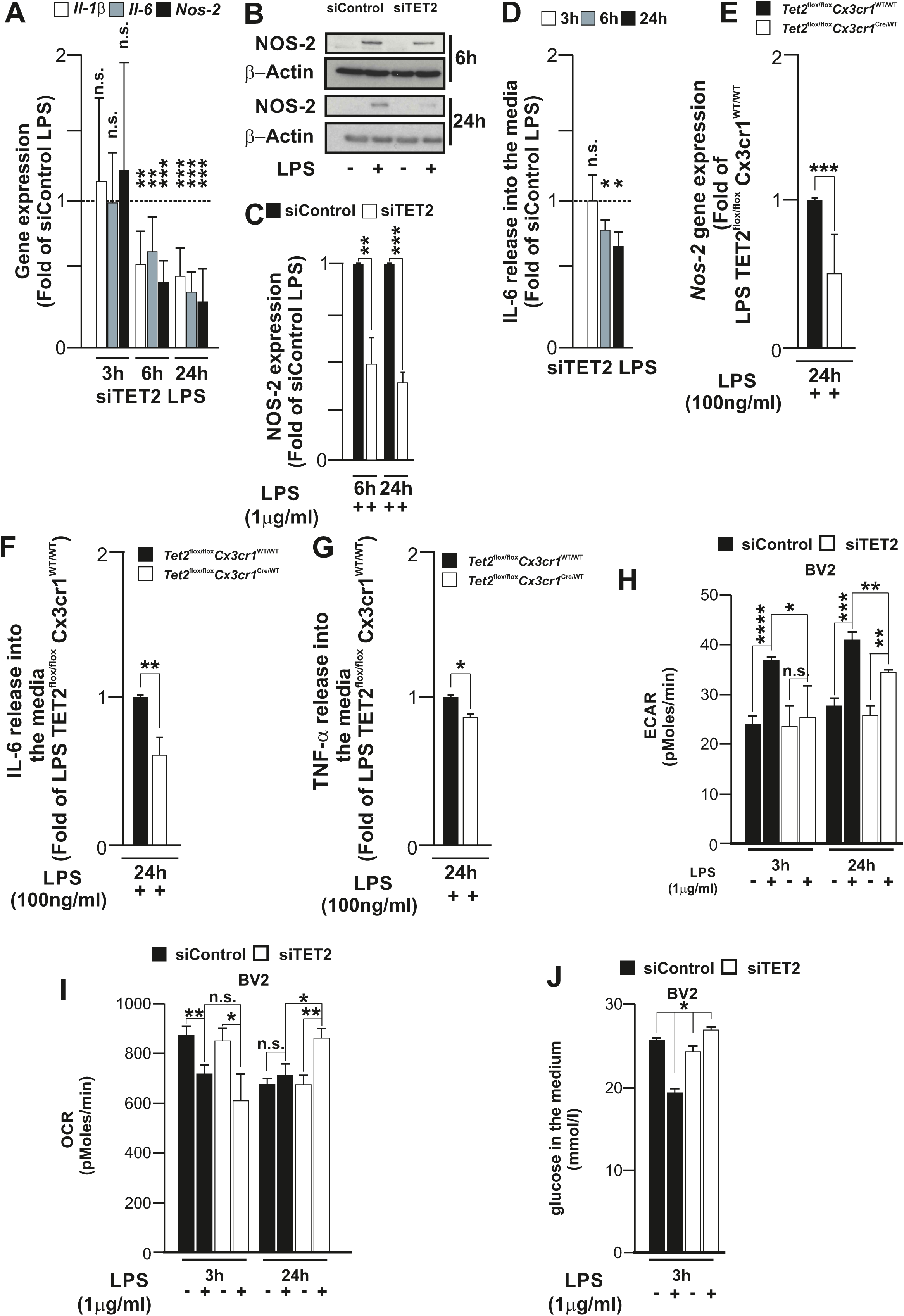
TET2 modulates LPS-induced changes in cellular metabolism and inflammatory response in microglia cells. Graph showing the gene expression of *Il-1β, Nos-2* and *Il-6* in LPS treated BV2 cells with or without TET2 gene knockdown at different time points (A). Representative immunoblot (B) and quantification (C) of NOS-2 protein at 6 h and 24 h LPS treatment in BV2 cells transfected with siRNA Control and siRNA *Tet2*. Quantification of IL-6 release into the media upon LPS treatment at different times (3, 6 and 24h) (D). Analysis of *Nos-2* gene expression in primary microglia cells after 24h treatment with LPS (E). Histograms showing the effect of TET2 gene knockdown over IL-6 (F) and TNF-α (G) in LPS treated primary microglia cells. Histograms showing the extracellular acidification rate (ECAR) (H), oxygen consumption rate (OCR) (I) and concentration of glucose (J) in the media in BV2 cells treated at different times with LPS. Data shown are mean ± s.d. of five (A) and three (C) independent experiments. Data shown in D are mean ± s.e.m of nine (3h), six (6h) and eight (24h) independent experiments. Data shown in E are mean ± s.d of seven independent experiments. Data shown in F and G are mean ± s.e.m of 9 (F) and 3 (G) independent experiments. Data shown in H, I and J are mean ± s.d of 3 independent experiments (H, I and J). Statistical analysis was performed using two-tailed Student’s t-test. \**P* < 0.05, \*\**P* < 0.01, \*\*\**P* < 0.001. See also Figure S5.

To validate our observations in primary microglia cell cultures, we crossed conditional *Tet2* floxed mice (*Tet2*^flox/flox^) with *Cx3cr1 Cre mice* (*Cx3cr1*^CreERT2/WT^) mice to enable the production of inducible microglia-specific deletion of *Tet2* (Figure S5A-C). We isolated primary microglia from both *Tet2*^flox/flox^*Cx3cr1* Cre-positive (henceforth referred to as *Tet2*^flox/flox^*Cx3cr1*^Cre/WT^) and negative mice (henceforth referred to as *Tet2*^flox/flox^*Cx3cr1*^WT/WT^), and treated cells with 4-OH-tamoxifen for 48h, producing a partial genomic deletion of *Tet2* (Figure S5A-B). Importantly, this led to decreased *Tet2* expression in *Tet2*^flox/flox^*Cx3cr1*^Cre/WT^ in basal conditions and complete abrogation of *Tet2* upregulation upon treatment with 100 ng/ml of LPS for 24 hours (Figure S5C). To analyze the effects of *Tet2* deletion over the inflammatory response, we compared the expression levels of different pro-inflammatory markers between primary *Tet2*^flox/flox^*Cx3cr1*^Cre/WT^ and *Tet2*^flox/flox^Cx3cr1^WT/WT^ in response to LPS. In accordance with our results from BV2 cells, *Nos-2* expression decreased by around 50% in *Tet2*^flox/flox^*Cx3cr1*^Cre/WT^ when compared with *Tet2*^flox/flox^*Cx3cr1*^WT/WT^ primary microglia 24h after LPS challenge (Figure 5E). We also analyzed the release of IL-6 and TNF-α into the media 24h after LPS treatment and observed a statistically significant inhibition of IL-6 release and TNF-α in primary microglia cell cultures supernatants (Figure 5F-G).

Collectively, our data suggest that, in microglial cells, TET2 is not involved in the resolution of the inflammatory response as reported in peripheral immune cells (Zhang et al., 2015). Instead and strikingly, microglial TET2 modulates the classical inflammatory response upon direct stimulus by LPS treatment. However, this time-dependent effect of TET2 on the LPS-induced expression of “classical” pro-inflammatory markers suggests an indirect effect. Therefore, we aimed to dissect the mechanisms that could affect the delayed expression of different inflammatory cytokines in activated microglia at different time points. TLR-4 stimulation induces a rapid and robust transcriptional response which involves genes that regulate metabolic reprogramming (Medzhitov and Horng, 2009). Treatment with LPS in macrophages, dendritic and microglia cells provokes a metabolic shift from oxidative phosphorylation (OXPHOS) towards aerobic glycolysis, a process required to quickly supply high energy demands of the inflammatory response (Galván-Peña and O’Neill, 2014),(Ruiz-García et al., 2011), (Ganeshan and Chawla, 2014),(Orihuela et al., 2016).

In pro-inflammatory (M1) macrophages, aerobic glycolysis is a consequence of glucose uptake and the conversion of pyruvate into lactate (Galván-Peña and O’Neill, 2014). Interestingly, two of the targets that were deregulated by *Tet2* knockdown in our RNAseq data were hexokinase 3 (*Hk3*) and 6-phosphofructo-2-kinase/fructose-2,6-biphosphatase 3 (*Pfkfb3*), both playing an important role during the aerobic glycolysis process (Galván-Peña and O’Neill, 2014)(Ruiz-García et al., 2011). We therefore asked if TET2 was involved in the early stages of the metabolic reprogramming induced by LPS. In BV2 cells, we measured the extracellular acidification rate (ECAR) as an indicator of lactate production and mitochondria oxygen consumption rate (OCR) as an indicator of the mitochondrial energy production in siControl and siTET2 BV2 cells with and without LPS at different time points (Figure 5H-I). A functional bioenergetics profile of siControl, siTET2 cells with and without LPS treatment in response to sequential treatment with oligomycin, FCCP and rotenone/antimycin A was carried out (Figure S5D-E). Our results show reduced lactate production after *Tet2* knockdown at 3 and 24h of LPS treatment at basal conditions (Figure 5H) and after oligomycin treatment, indicating reduced glycolysis (Figure S5D). We then asked whether this decrease in lactate formation at 3h was correlated with a decrease in the glucose consumption. We analyzed the extra-cellular glucose levels after 3h LPS treatment and found that siControl LPS treated cells consume glucose from the media, but that LPS-treated siTET2 cells show a substantial reduction in the glucose uptake (Figure 5J). Because *Tet2* knockdown strongly reduced LPS-induced glycolysis, and because microglia have a metabolic dependence on glycolysis (Vilalta and Brown, 2014), we tested whether inhibition of glycolysis affected the inflammatory response. In agreement with our previous data, inhibition of hexokinase activity by using 2-deoxy-D-glucose (2-DG) inhibited the inflammatory response measured as NOS-2 expression in BV2 at 6h without inducing cell death (Vilalta and Brown, 2014), (Tannahill et al., 2013) (Figure S5F-G). These results suggest that TET2 regulation of the inflammatory response might be mediated by the early changes in glycolysis induced by TET2. Indeed, TET2 knockdown reduced the basal and maximal oxygen consumption of the cells after 3h LPS treatment (Figure 5I and Figure S5E), but after 24h LPS treatment, the cellular oxygen consumption levels increase (Figure 5I). This indicates that TET2 mediates the substantial metabolic reprogramming of the cells induced by LPS, including an early rise in glycolytic and mitochondrial energy production, followed by a fall in mitochondrial energy production potentially mediated by the well-known inhibition of mitochondria by NO from NOS-2 (Bal-Price et al., 2002) (Kelly and O’Neill, 2015) (Doulias et al., 2013).

### Effect of TET2 depletion in microglia cells in vivo

Our *in vitro* results using primary microglia cells and BV2 cell line suggest that TET2 is necessary for a full proinflammatory response. These results contrast with findings in peripheral macrophages after intraperitoneal injection *in vivo* (Zhang et al., 2015). We therefore assessed the effect of microglial TET2 depletion *in vivo*, using an inflammatory model based on intraperitoneal injection of LPS, in our *Tet2*^flox/flox^*Cx3cr1* ^Cre/WT^ and *Tet2*^flox/flox^Cx3cr1^WT/WT^ mice (Figure S6A). It has been demonstrated that this *in vivo* model induces a well-defined microglial proinflammatory phenotype different to the recently characterized molecular signature of disease-associated microglia (Bodea et al., 2014; Krasemann et al., 2017).

In our conditional inducible KO mouse model, we achieved around 40% decrease in the expression of TET2 at the protein level within microglial cells (Figure S6B-C). We first assessed the effect of TET2 depletion in microglia cells after intraperitoneal injection of LPS in the substantia nigra (SN), in *Tet2*^flox/flox^*Cx3cr1*^Cre/WT^ and *Tet2*^flox/flox^Cx3cr1^WT/WT^ mice (Figure 6A-G). We observed that depletion of microglial TET2 failed to affect microglia cell density at physiological conditions. However, upon treatment with LPS, we observed a decrease in the proliferation rate in microglia lacking TET2 (Figure 6A-B). This result is supported by our RNA-seq in BV2 cells, where we showed that several genes involved in cell cycle regulation were under control of TET2. Microglia activation is well known to be associated to prominent morphological alterations. In SN, while homeostatic microglia are highly ramified cells, upon activation, microglia increase cell body and Iba1 expression along with thickening of processes to end in complete retraction of cytoplasmic processes to acquire an amoeboid morphology. In response to repeated systemic LPS injections, a massive appearance of microglia exhibiting typical morphological features of activation was found 24h after LPS challenge (Figure 6A). This observation prompted us to perform a detailed analysis of microglia based on morphological features of homeostatic microglia and three well-defined states of microglia activation as shown in Figure 6C. TET2 deletion did not alter the density of homeostatic microglia in healthy unlesioned brain (Figure 6C). In response to LPS, *Tet2*^flox/flox^*Cx3cr1*^WT/WT^ mice showed very low presence of homeostatic microglia, with very robust increases of activated microglial cells (Figure 6C). *Tet2*^flox/flox^*Cx3cr1* ^Cre/WT^ mice showed significantly lower degree of microglia activation, affecting the number of reactive microglia cells (Figure 6C). A typical early feature of activated microglia is cell proliferation (Mathys et al., 2017), hence an effect of TET2 in cell cycle regulation cannot be discarded in response to proinflammatory challenge in line with our RNA-seq data. We next dissected out the SN and extracted mRNA to measure the expression of different cytokines. TET2 knock-down in microglia cells failed to affect the expression of *Il-1β, Tnf-*α and *Nos-2* in response to systemic LPS (Figure S6D). It is, however, important to note that in this *in vivo* model microglia are not directly activated by LPS binding to TLR-4, but by different factors released by activated peripheral immune cells (Chen et al., 2012). We therefore also analyzed the effect of TET2 deletion on genes that become highly expressed after intraperitoneal injection with LPS. Based on publicly available data in this model (Krasemann et al., 2017), we focused on the expression of the two highest induced genes upon repeated systemic LPS injections, *Ptgs2* and *Cybb* (also known as *Cox-2 and Nox-2* respectively), where both genes are considered as proinflammatory mediators (Alhadidi and Shah, 2018; Benusa et al., 2017). While no difference in expression was found with *Nox-2* expression in LPS-treated TET2 depleted mice (Figure 6D), there was a very strong inhibition of the LPS-induced COX2 expression in TET2 depleted microglia cells both at mRNA and protein levels (Figure 6E-G). Remarkably, these results were obtained with a partial repression in TET2 expression of 40%. Therefore, we cannot exclude the possibility that a higher degree of repression of TET2 may have also a wider effect affecting the expression of *Nox-2* as well.

**Figure 6:**
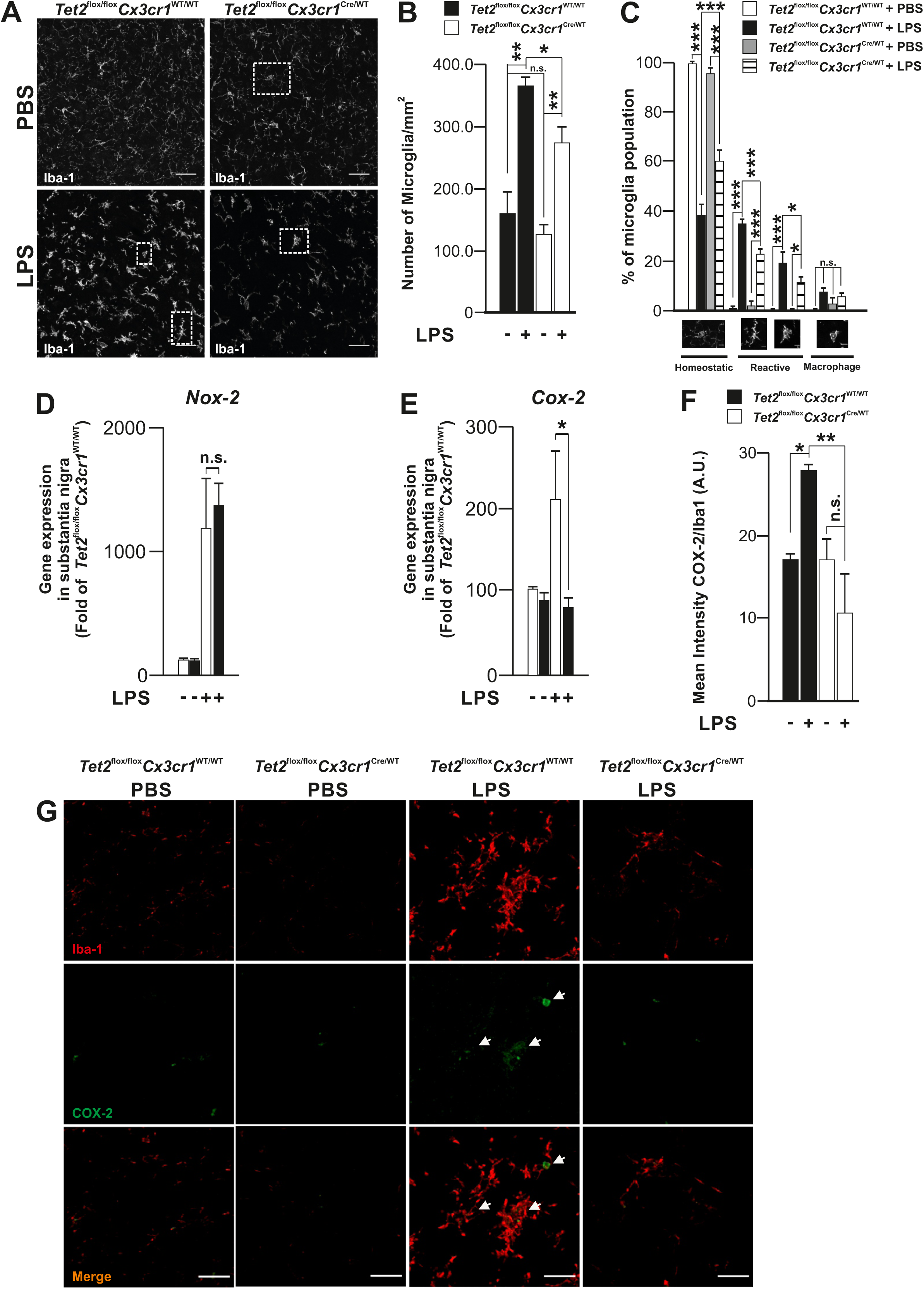
Abrogation of *Tet2* in microglia cells decreases the LPS-induced immune response in vivo. Iba-1 immunostaining in substantia nigra of *Tet2*^flox/flox^*Cx3cr1* ^Cre/WT^ and *Tet2*^flox/flox^Cx3cr1^WT/WT^ mice treated for four days either with LPS or vehicle (PBS) and sacrificed 24h later (A) and analysis of microglia cell numbers (B) and activation status based on morphology (C). qRT-PCR analysis of *Nox-2* and *Cox-2* in *Tet2*^flox/flox^*Cx3cr1* ^Cre/WT^ and *Tet2*^flox/flox^Cx3cr1^WT/WT^ mice treated for four days either with LPS or vehicle (PBS) and sacrificed 24h later (D and E). Quantification of Cox-2 staining in Iba-1 positive cells in the same type of mice (F and G). Data represented as mean ± s.e.m. The results correspond to three independent experiments in panels B and C. In panels D and E the number of independent experiments is equal to three for *Tet2*^flox/flox^*Cx3cr1*^WT/WT^ + LPS and seven for *Tet2*^flox/flox^*Cx3cr1*^Cre/WT^ + LPS. In panel G all treatments are 3 independent experiments except *Tet2*^flox/flox^*Cx3cr1*^Cre/WT^ + LPS which is four independent experiments. Statistical analysis was performed using one-way ANOVA with a Student-Newman-Keuls Method post hoc test (C, G) or two-tailed Student’s t-test (D, E). \**P* < 0.05, \*\**P* < 0.01, \*\*\**P* < 0.01. See also Figure S6.

Altogether these results show that TET2 plays a pro-inflammatory role in microglia (opposite to peripheral immune cells) and highlight fundamental differences in the role of TET2 in the inflammatory response between microglia and peripheral immune cells. This difference in behavior with peripheral macrophages and dendritic cells has been already described in injured CNS conditions where, for instance, monocyte-derived macrophages often take a more CNS “repair/supportive” phenotype than resident microglia under ischemic conditions (London et al., 2013; Zarruk et al., 2018).

### Microglial TET2 expression during the neurodegenerative process

We and others (Janova et al., 2016), have observed that *Tet2* expression is induced under inflammatory conditions mediated by different TLR agonists and fibronectin (Figure S1A-C). Since microglia play a fundamental role in the demise of the neuronal population in several neurodegenerative diseases, we tested whether *Tet2* expression could also be affected by different pathological protein aggregates seen in a number of neurodegenerative diseases typically associated to neuroinflammation (Ugalde et al., 2016). For instance, α-synuclein aggregates induce microglial activation (Boza-Serrano et al., 2014) in a TLR-4 dependent manner (Fellner et al., 2013). Similarly, the fibrillar and oligomeric Aβ-amyloid-induced neuroinflammatory response has been linked to TLR4 as well (Reed-Geaghan et al., 2009; Wang et al., 2018). Therefore, we challenged our BV2 cells with α-synuclein aggregates and Aβ-amyloid oligomers for 6h and observed an increase of *Tet2* expression similar to that observed with LPS (Figure S7A). To test whether microglial TET2 is upregulated *in vivo*, we analyzed TET2 expression in an AD mouse model. We used the 18 month-old 5xFAD mouse model, which mirrors the main features of AD through five mutations linked to familial forms of AD, and develops extracellular amyloid plaques typically associated with clusters of highly reactive microglia. This model recapitulates in a few months the main features of AD (Oakley et al., 2006). Strikingly, we observed in hippocampus that plaque-associated microglia displayed increased TET2 expression when compared to homeostatic microglia located further from the plaques (homeostatic microglia) (Figure 7A-B and Figure S7B).

**Figure 7:**
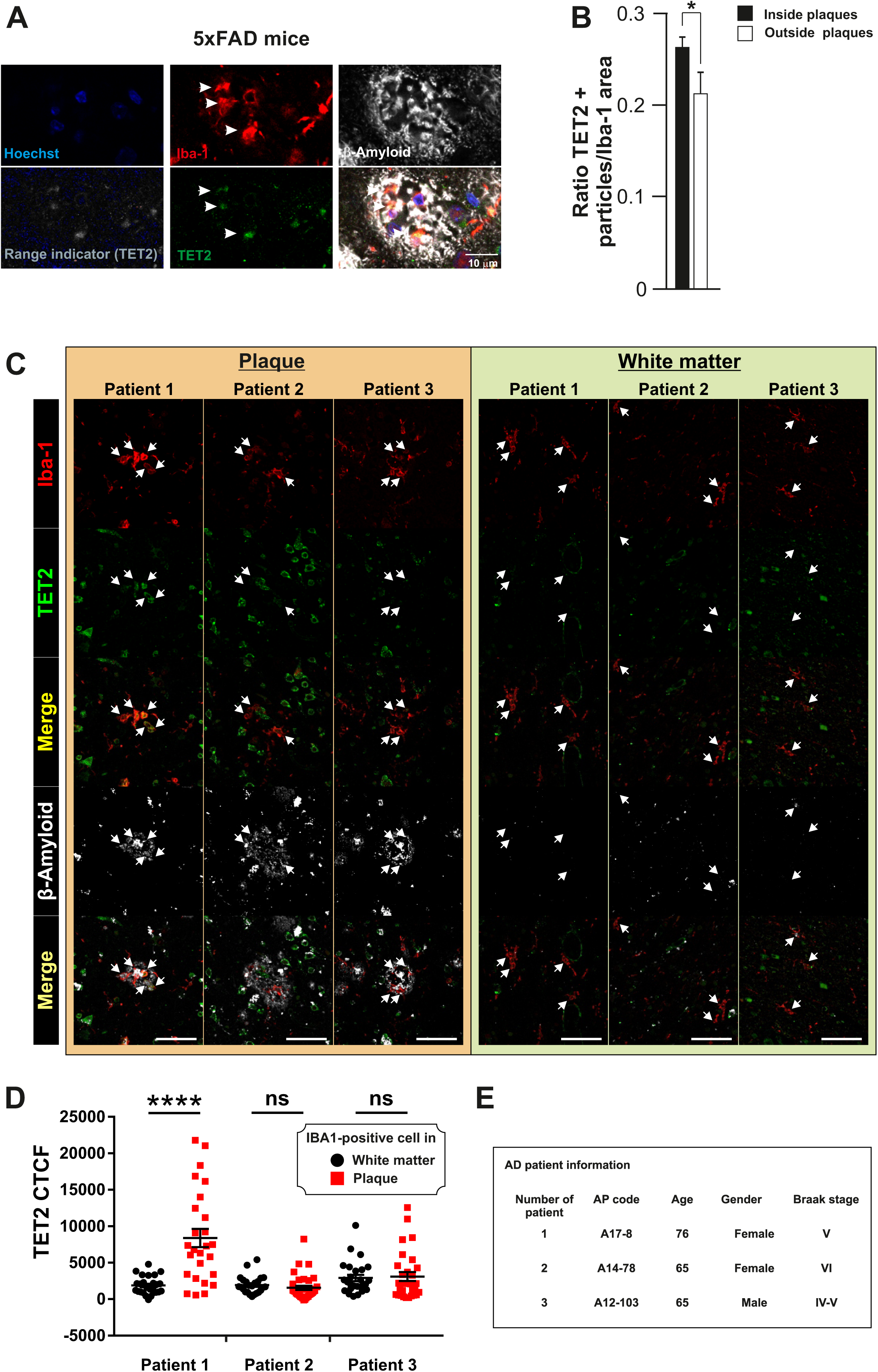
Microglial TET2 expression in 5xFAD mice and Alzheimer’s disease brain tissue. Colocalization analysis of TET2, Iba-1 and beta amyloid (β-plaques) in the hippocampus of 18 month-old 5xFAD mice (A, B) and three AD patients (C, D). Panel E shows details of age, gender and Braak stage. Data shown in B is mean ± s.e.m of 3 independent experiments. Arrowheads indicate microglial cells. CTCF stands for “corrected total cell fluorescence” See also Figure S7. \**P* < 0.05

Finally, in order to test the clinical relevance of these results, we analyzed TET2 expression in human microglia cells in post-mortem temporal cortex tissue from three AD patients (Figure 7C-E). We measured the fluorescent intensity of TET2 expression in Iba-1-positive microglial cells associated to Aβ plaques. These values were compared to TET2 expression in Iba-1-positive microglial cells located in the white matter, and therefore not associated to plaques. Our analysis showed that in one patient (patient 1), there is significant upregulation of TET2 in plaque-associated microglia, while no statistical significance was observed in the other two patients (Figure 7D). The discrepancy in the human samples cannot be directly linked to either age, gender or Braak stage of the patients (Figure 7E), and more thorough studies are needed to clarify these differences. Of note, plaques in AD patients are very heterogeneous (unlike 5xFAD mice) which may explain the difference in microglial TET2 among the three patients. Also, many TET2-positive cells did not colocalize with Iba-1-positive cells (Figure 7D), Based on their morphology, these TET2-positive/Iba-1-negative cells could be neurons, which agrees with previous studies showing increased expression TET2 expression in neurons (Mi et al., 2015)(Svetlana Dzitoyeva, Hu Chen, 2008). We observed that the ratio of Iba-1+/TET2+ cells vary within the three patients. While all the patients present some Iba-1 positive cells expressing TET2, the numbers vary greatly within the three patients. The Iba-1+/TET2+ ratio is relatively high in patient number 1, but on the other hand, patients number 2 and 3 present a lower ratio (Figure 7D). Importantly, similar to the result obtained in the 5xFAD mouse model, in one of the patients microglial TET2 expression is highly expressed in the plaque-associated microglia (DAM microglia), while microglia localized away from the plaques (homeostatic) showed little or no induction of TET2 expression (Figure 7A-D). This differential response of TET2 expression in microglia cells depending on the distance to the β-plaque suggests that β-amyloid might also be a direct or indirect inducer of TET2 expression over time.

Altogether, these results suggest that TET2 could play an important role in the neuroinflammatory response driven by microglia, among others, in AD and PD, although more comprehensive studies are necessary.

## Discussion

Epigenetic mechanisms have been proposed to regulate distinct aspects of the inflammatory response in different immune cell types. Recent reports have shown that TET2 plays important roles during the inflammatory response in different peripheral immune cells (Ichiyama et al., 2015)(Tsagaratou et al., 2016)(Zhang et al., 2015)(Cull et al., 2017). In this study, we demonstrate that TET2 is early upregulated in microglia in response to TLR activation in a NF-κB-dependent transcriptional process. We provide strong evidence that TET2 drives early expression of multiple genes associated to the immune system including the Type I IFN response upon TLR stimulation, an effect that was largely independent from its oxidative activity. Notably, TET2 played central roles in driving both the classical proinflammatory response and metabolic reprogramming that take place during the TLR-dependent microglia activation. The potential involvement of TET2 in the recent characterized disease-associated microglia (Keren Shaul et al., 2017; Krasemann et al., 2017) is deduced from analysis of both AD transgenic mice and human AD tissues in which TET2 was highly upregulated specifically in amyloid plaque-associated microglia.

We found that TET2 regulates primarily genes involved in the innate immune response and more specifically genes related to the TLR induced-Type I IFN response (e.g., *Stat1, Stat3, Irf1, Irf7*) (Noppert et al., 2007)(Luu et al., 2014). Traditionally, Type I IFN response was considered solely for defense against viral and bacterial infections (Stifter and Feng, 2015)(Kovarik et al., 2016). However, in the brain, and under sterile inflammatory conditions, there are increasing numbers of reports showing activation of the Type I IFN response in ischemia (McDonough et al., 2017), spinal cord injury (Impellizzeri et al., 2015) and in two different AD mouse models, the 5xFAD (Landel et al., 2014) and APP/PS1 (Taylor et al., 2014), as well as in AD patients (Taylor et al., 2014). In fact, using single-cell RNA sequencing of microglia at different stages in a severe neurodegeneration model (CK-p25 mouse model) for AD (Mathys et al., 2017), the authors described that “late-stage” activated microglia is characterized by the expression of many Type I and Type II Interferon response genes. Incidentally, in the same study, and similar to our data, gene ontology analysis show an enrichment in the expression of genes involved in the control of cell cycle in “early-stage” activated microglia. These results highlight the similarity of the TET2 regulated inflammatory response in our experimental conditions with other neurodegenerative models.

Our RNA-seq data show that TET2 regulates the early (3h) transcriptional response to LPS but without an immediate effect on traditional markers for the inflammatory response. Indeed, 6-24h after treatment, we observed a repression in the expression of the classic markers *Il-1β, Il-6* and *Nos-2* in *Tet2* knockdown BV2 cells. This effect was confirmed in primary microglia cells where *Tet2* was deleted. This result raises the importance of TET2 in governing proinflammatory activation in microglia. Notably, TET2 has been shown to specifically repress *Il-6* transcription at late phase in LPS-treated macrophages to terminate the inflammatory response (Zhang et al., 2015). Using the same *in vivo* model used by Zhang and colleagues, we observe that TET2 is also upregulated in microglia but TET2 is necessary for a full proinflammatory response. This effect differs from the result reported here and highlights fundamental differences in the regulation of the innate immune responses between peripheral immune cells (and specifically monocyte-derived macrophages) and microglia (Burm et al., 2015) (London et al., 2013; Zarruk et al., 2018), suggesting that the action of TET2 is highly context-specific.

How does TET2 influence the late phase of the classical inflammatory response? There is growing acknowledgement of the role that the Type I IFN response plays over the TLR-induced classical inflammatory response (Luu et al., 2014)(Taylor et al., 2014). Of note, one of the TET2-regulated genes that forms part of this response is the IFN-induced protein with tetratricopeptide repeats 2 (IFIT2), which has been shown to play a role in the amplification of the secretion of TNF-α and IL-6 *in vivo* in a LPS-induced endotoxin shock model (Siegfried et al., 2013). Additionally, TET2 regulates a number of small IFN-induced GTPases, and in particular guanylate-binding proteins (GBPS). It has been reported that GBPS are required for the full activation of the non-canonical caspase-11 inflammasome activation and for the secretion of IL-1β during infections with vacuolar Gram-negative bacteria, (Meunier et al., 2014). GBP2 and GBP3 appear in our RNA-seq analysis as genes under TET2 control upon LPS treatment, which could potentially affect the inflammatory response.

Our data also suggest that TET2 could regulate the classical inflammatory response through modulation of LPS-induced changes in metabolism. LPS causes a major shift in the metabolism of macrophages, dendritic cells, neutrophils and microglia from oxidative phosphorylation towards aerobic glycolysis, similar to what happens in tumor cells in a process known as the Warburg effect (Rodríguez-Prados et al., 2010) (Orihuela et al., 2016). We found that TET2 depletion leads to reduced glucose consumption and lactate production, an effect that precedes the reduction in classic inflammatory markers. Interestingly, two genes known to regulate glycolysis (*Hk3 and Pfkfb3*) are regulated by TET2 upon LPS treatment. Hexokinase 3 catalyzes the first committed step of glycolysis (Nishizawa et al., 2014), while PFKFB3 catalyzes both the synthesis and degradation of fructose-2,6-bisphosphate (F2,6BP), a regulatory molecule that controls glycolysis in eukaryotes. Direct inhibition of glycolysis (using 2-D-Deoxyglucose to inhibit hexokinase activity) prevented LPS induction of NOS-2, suggesting that the TET2 regulation of glycolysis may mediate the LPS induced inflammatory response.

We also found that microglial TET2 is upregulated *in vivo* in two-well defined models of microglia polarization; in a neuroinflammatory mouse model induced by repeated intraperitoneal injections of LPS (Krasemann et al., 2017; Bodea et al., 2014), and in AD, including human patients and transgenic mice (5xFAD). In AD, upregulated TET2 was restricted to plaque-associated microglia, which has been largely associated to AD pathogenesis. AD and PD are characterized by the accumulation of aggregated proteins; i.e. intracellular α-synuclein in PD and extracellular amyloid β in AD, whose immunogenic properties have been well documented. In fact, these aggregated forms play leading roles in driving main pathogenic events in these neurodegenerative diseases with the active involvement of highly activated microglia, more recently referred to as disease-associated microglia (Krasemann et al., 2017) (Keren-Shaul et al., 2017) (Mathys et al., 2017). In human AD and in transgenic mice, disease-associated microglia are strictly confined to amyloid β plaques (plaque-associated microglia (Keren-Shaul et al., 2017)). It is noteworthy to mention that in different single cell RNAseq of microglia in different AD models, the expression of TET2 was found not to be upregulated (Krasemann et al., 2017) (Keren-Shaul et al., 2017) (Mathys et al., 2017). This is interesting because our analysis using an antibody against TET2 (validated in Knockout mice) in temporal cortex of human AD and hippocampus of 5xFAD mice demonstrated the unequivocal upregulation of TET2 in disease plaque-related microglia but not in homeostatic microglia. This suggests that increased levels of TET2 protein present in microglia cells might result from a post-translational mechanism (for instance a decrease in the rate of protein degradation which allows an increase of protein content).

Our results strongly support the idea that TET2 could drive the proinflammatory activation of microglia and induction of metabolic reprogramming upon inflammatory stimulus. Keeping in mind the chronic nature of inflammation in neurodegenerative diseases, an active role of TET2 in the switch from homeostatic to disease-associated microglia is anticipated. In the future, TET2 may become a potential drug target to control exacerbated neuroinflammatory response in neurodegenerative diseases.

## Methods

### Cell Lines, Transfection, Reagents and accession number of RNAseq data

Human CHME3 and murine microglial BV2 cell line were cultured as described in (Bocchini et al., 1992). Briefly, the cells were maintained in 10% fetal calf serum (FCS) in DMEM and reduced to 5% FCS during the experiments. Transfection of BV2 cells was carried out using Lipofectamine 3000 (Invitrogen) following the manufacturer’s instructions. A complete list of reagents, siRNA, primers, antibodies and patient information are provided in the Key resources tables. The accession number of our RNAseq data is GSE105155.

### Generation of microglia specific Tet2-deficient mice

All experiments conducted with animals were previously approved by the different Ethical Committee for Experimental Research from University of Seville and University of Cambridge and fulfilled the requirements for experimental animal research in accordance with in accordance with the U.K. Animals (Scientific Procedures) Act (1986) and the Guidelines of the European Union Council (86/609/EU) and the Spanish and UK regulations (BOE 34/11370– 421, 2013) for the use of laboratory animals.

*Tet2*^flox/flox^ C57BL/6 mice with the *Tet2* allele floxed at exon 3 (Jackson Laboratories, B6;129S-Tet2tm1.1Iaai/J) and C57BL/6 mice containing a Cre recombinase under the control of *Cx3cr1* promoter and enhancer elements (Jackson Laboratories, B6.129P2(Cg)-Cx3cr1tm2.1(cre/ERT)Litt/WganJ), were crossed to generate *Tet2*^flox/flox^;*Cx3Cr1*^Cre/WT^ (experimental mice) and *Tet2*^flox/flox^;*Cx3Cr1*^WT/WT^ (control mice).

Animals were housed under a 12h light/dark cycle with free access to food and water. The genotype of *Tet2*^flox/flox^;*Cx3Cr1*^WT/WT^ and *Tet2*^flox/flox^;*Cx3Cr1*^Cre/WT^ mice was determined by analysis of DNA extracted from the fingers using a QuickExtract™ (Epicentre) and amplified with MyTaq™ Red DNA Polymerase (Bioline). The deletion of the *Tet2* gene was determined by analysis of DNA extracted from isolated primary microglia using a QuickExtract™ (Epicentre) and amplified with MyTaq™ Red DNA Polymerase (Bioline).

The PCR consisted of 94 °C for 1min, then 35 cycles with denaturation at 95 °C for 15 s, annealing at 58 °C for 15 s, and elongation at 72 °C for 10 s. The primer sequences used were obtained from a previous study (Moran-crusio et al., 2011).

### Data and Software Availability

The accession number for the RNA-seq data reported in this paper is NCBI Gene Expression Omnibus: GSE105155.

## Author contributions

M.A.B and M.R.B. designed the study. A.C-J,J.L.V, I.G-D, J.G-R, J.C. F-M and R.R performed all the experiments performed with the *Tet2*^flox/flox^; *Cx3Cr1*^Cre/wt^ and *Tet2*^flox/flox^; *Cx3Cr1*^wt/wt^ mice. Ö.D performed the ChIP analysis for NF-κB p65, and H3K27ac and H3K4me1 abs. Ö.D and M.A.B performed the ChIP analysis for TET2 ab. M.V. N-C determined the ECAR and OCR index and glucose consumption in BV2 cells. K.B-S performed the oxBS sequencing experiments. V.S, and X.S. performed the staining and analysis of TET2 expression in AD patients. R.A and P.H. performed the mass spectrometry analysis for global levels of 5mC and 5hmC. A.V and G.C.B performed the analysis of TET2 in rat postnatal primary microglia. P.K.Y performed the analysis of TET2 in mouse adult primary microglia. M. C performed the analysis of TET2 in the human microglia cell line CHME3. R.R. performed the analysis of TET2 in the 5xFAD mice brain tissue and in *Tet2*^flox/flox^;*Cx3Cr1*^WT/WT^ and *Tet2*^flox/flox^;*Cx3Cr1*^Cre/WT^ injected with LPS. R.R also performed the confocal analysis of microglia morphology and COX-2 expression in *Tet2*^flox/flox^;*Cx3Cr1*^WT/WT^ & *Tet2*^flox/flox^;*Cx3Cr1*^Cre/WT^ injected with LPS. A.M.S.E. analyzed TET2/3 expression in BV2 cells after LTA treatment. K.T. and A.M. E-O. and P.S-H analyzed the expression levels of TET2 in BV2 upon α-synuclein and β-oligomer treatments. E.R. provided the AD patient tissue. I.G-D, J.G-R, J.C F-M performed the isolation of RNA from substantia nigra from *Tet2*^flox/flox^;*Cx3Cr1*^WT/WT^ & *Tet2*^flox/flox^;*Cx3Cr1*^Cre/WT^ injected with LPS and performed the qPCR analysis for the different cytokines and COX2 and NOX2. M.R.B performed the RNA-seq experiments and analyzed the data. M.A.B performed the qPCR analysis and immunoblots for the study of the inflammatory response in BV2 cells. J.L.V and B.J were involved in the study design. M.A.B. and M.R.B wrote the manuscript with all the authors. M.A.B., M.R.B and J.L.V share senior authorship and M.A.B and M.R.B share corresponding authorship in this paper. All authors discussed the results and commented on or edited the manuscript.

## Acknowledgments

We would like to thank to Dr. de la Rica (Queen Mary University of London) for technical support and Dr. Hornedo-Ortega (University of Seville) for providing us with the α-synuclein. A.C.-J. is supported by a pre-doctoral fellowship from Spanish Ministerio de Educación, Cultura y Deporte. Ö.D. has received funding from the People Programme (Marie Curie Actions) of the European Union’s Seventh Framework Programme (FP7/2007-2013) under REA grant agreement no 608765. M.V.N-C received a Children with Cancer UK fellowship (14-178). This work has been supported by grants from Blizard Research Theme grants Queen Mary University of London, Blizard Institute, Spanish Ministerio SAF2015–64171R (MINECO/FEDER, UE), Swedish Research Council, ERA-NET TracInflam, Swedish Brain Foundation and Karolinska Institutet Foundations. M.R.B. is a Sir Henry Dale Fellow (101225/Z/13/Z), jointly funded by the Wellcome Trust and the Royal Society. M.A.B has been funded by the Barts Charity Centre for Trauma Sciences (C4TS) and the Wellcome Trust Institutional Strategic Support Fund (ISSF) and the Spanish Ministry of Economy and Competitivity (Programa Ramón y Cajal: RYC-2017-21804). The authors declare no conflict of interest.

