## Supplemental Figures 1 to 7 for "TET2 regulates the neuroinflammatory response in microglia"

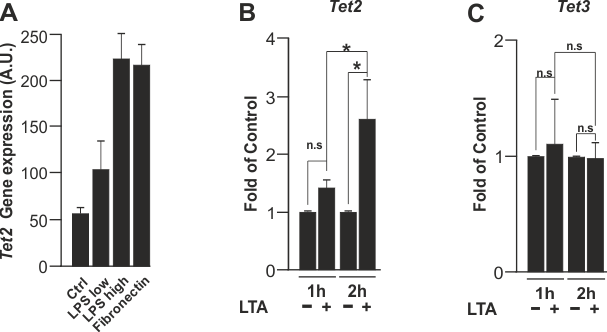

**Figure S1 (related for Figure 1): TLR2 and TLR4 activation induces *Tet2* upregulation in microglial cells.**

*Tet2* expression after different proinflammatory stimuli in primary postnatal murine microglia generated from the RNAseq data obtained from (Janova et al., 2016) (A). *Tet2* (B) and *Tet3* (C) gene expression after 1h and 2h treatment with 50 μg/ml lipoteichoic acid (LTA) in BV2 cells. Data are represented as mean ± s.d. of three independent experiments (A, B, C), using one-way ANOVA with Scheffe correction (B, C). **P* < 0.05

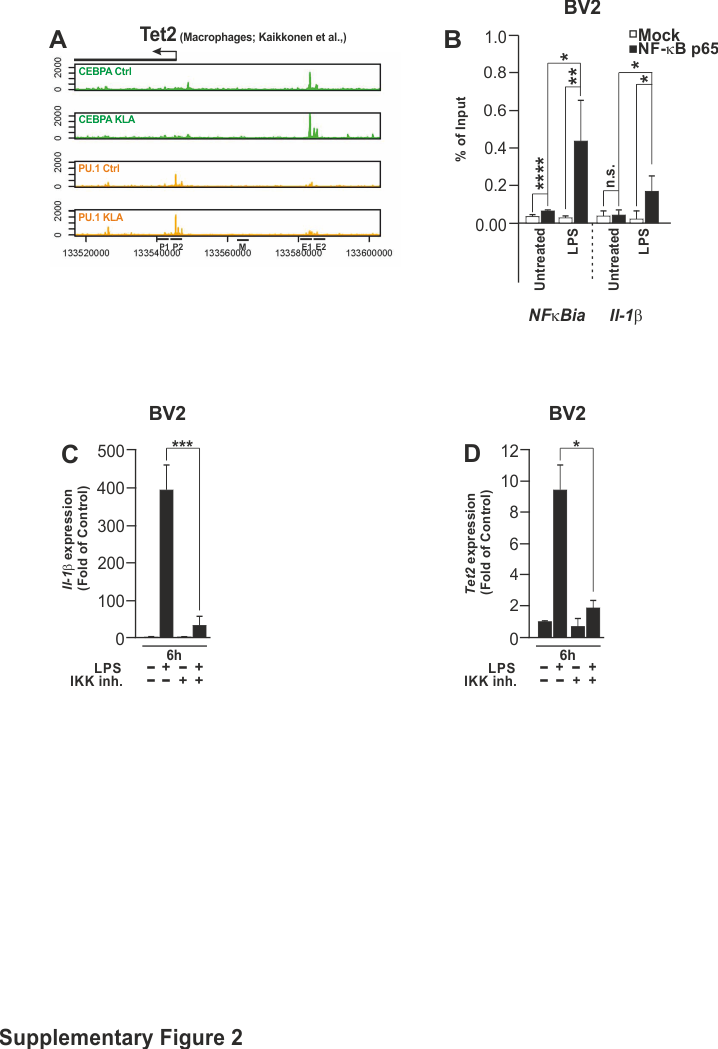

**Figure S2 (related to Figure 2): ChIP-seq profile of *Tet2* promoter after TLR4 activation.**

Analysis of data from the study published in (Kaikkonen et al., 2013), covering the *Tet2* promoter and upstream regions for CEBPA and PU.1 transcription factors in peripheral macrophages treated with KLA (an agonist for TLR-4) (A). *NFκBia* and *Il-1β* were used as positive controls of genes under p65 transcriptional control upon LPS treatment in BV2 microglia cells (B). Measurement of LPS-induced *Il-1β* (C) and *Tet2* expression (D) with and without pre-treatment of wedelactone (30 μM) for 1 h. LPS-induced *Il-1β* was used as a positive control for a gene regulated through NF-κB p65 in BV2 microglia cells. Data represented in B are representative replicate of two biological replicates. Data represented in C and D are represented as mean ± s.d from three independent experiments. Statistical analysis was performed using two-tailed Students t-test (C, D). **P* < 0.05, ****P* < 0.001

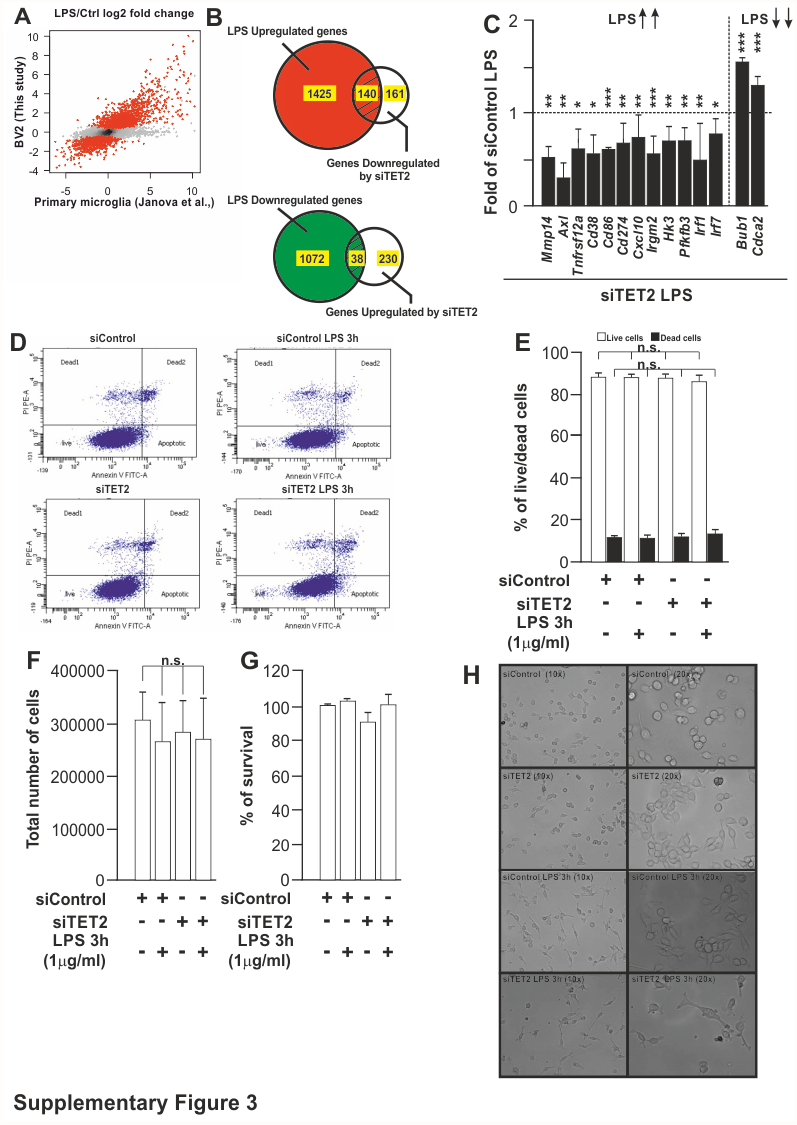

**Figure S3 (related to Figure 3): Validation of several targets obtained from the RNAseq. data**

Scatter plots comparing our RNA-seq data generated in BV2 cells (Y axis) with the RNA-seq data obtained from a published study using primary postnatal microglia (Janova et al., 2016) (X axis). The log2 fold-change in gene expression upon LPS treatment is plotted. Red data points represent genes that are differentially expressed in BV2 cells when comparing LPS-treated cells with untreated controls (A). Venn diagrams generated from our RNAseq data in BV2 representing the number of genes affected by LPS treatment and siRNA TET2 treatment. Validation of the effect of TET2 knockdown on gene expression on selected gene targets obtained in our RNA-seq (C). Dot plot (D) and bar chart (E) of the analysis of Annexin V/PI staining in BV2 cells transfected with siControl and siTET2 for 48 hours and treated with LPS (1μg/ml) for 3 hours. Analysis of total number of cells (F) and % of survival (G) in staining in BV2 cells transfected with siControl and siTET2 for 48 hours and treated with LPS (1μg/ml) for 3 hours. Bright field photographs of BV2 cells transfected with siControl and siTET2 for 48 hours and treated with LPS (1μg/ml) for 3 hours (H). Data shown are represented as mean ± s.d. from three (C-G)to six (C) independent experiments. Two-tailed Student's t-test. *P < 0.05, **P < 0.01, ***P < 0.001

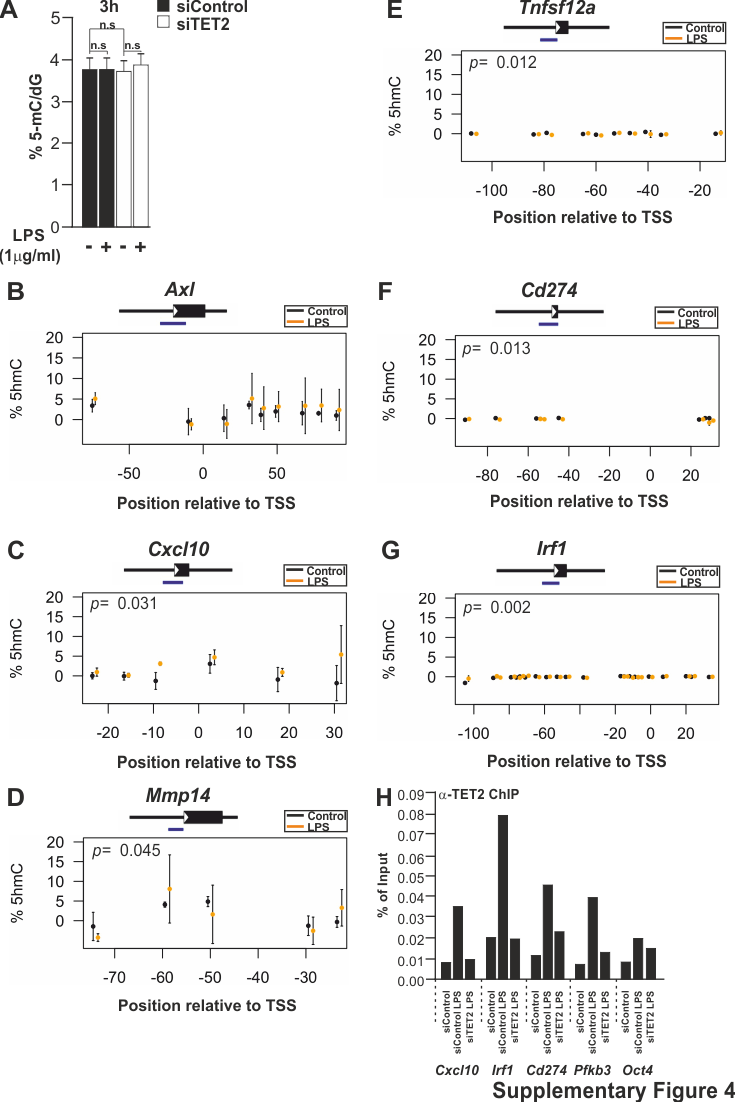

**Figure S4 (related to Figure 4): Oxidative bisulfite (oxBS) sequencing of different genes affected by LPS treatment.**

Global DNA methylation analysis in transfected microglia (siControl and siTet2) with and without 3h treatment with LPS (1μg/ml). Data shown represent the mean ± s.d from three independent experiments and statistics were performed using two-tailed Student’s t-test (A). Quantification by oxBS-seq of 5-hmC levels after 3h LPS treatment of different target genes (blue bars indicate the position of the analysed amplicons) (B-G). TET2 ChIP of *Cxcl10 , Irf1 Cd274, Pfkb3* and *Oct4* (used here as a negative control) genes after 3h treatment with LPS (1μg/ml) and including a siRNA Tet2 LPS control (H). Data shown represent the mean of two technical replicates. Statistics were performed using two-way ANOVA with a Tuckey post-hoc test tacking into account all CpGs. We include the *p* value for genes with a statistically significant difference in the graphs (B-G). Data representative of one technical replicate in H.

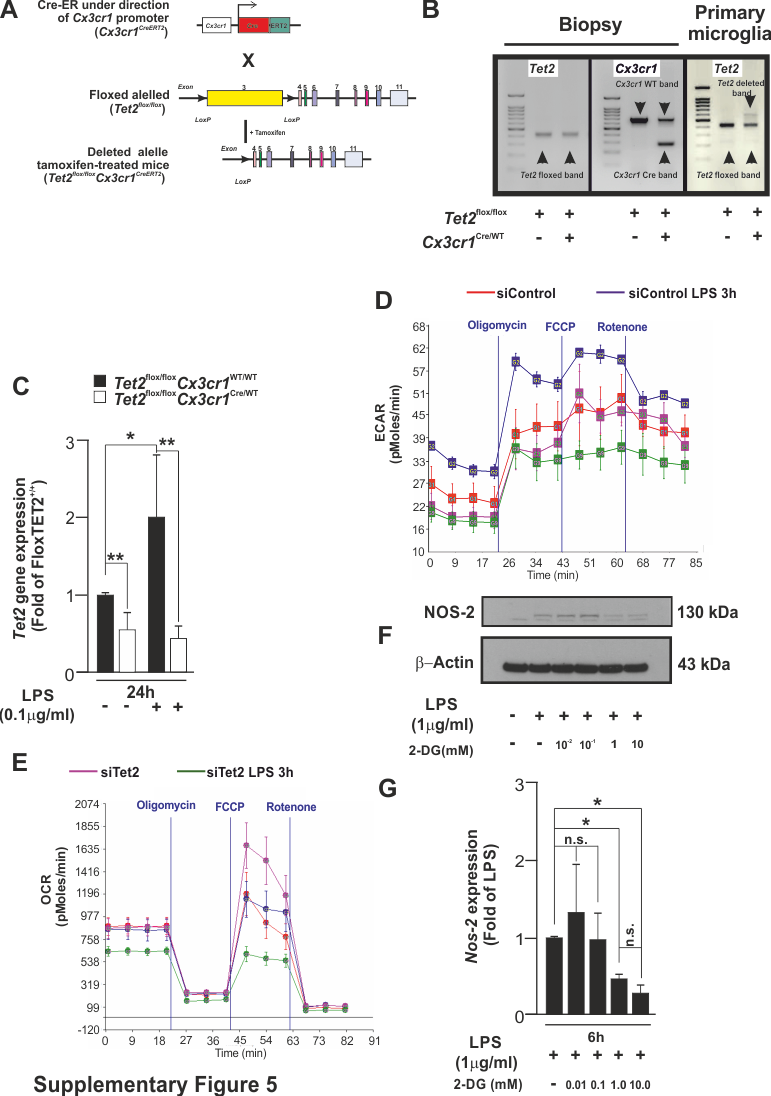

**Figure S5 (related to Figure 5): Representative example of ECAR and OCR graphs and the effect of 2-deoxy-D-glucose on the inflammatory response induced by LPS**

A schematic diagram depicting the mechanism of how treatment with 4-hydroxitamoxifen promotes the excision of Tet2 allele 3 by using the Cre-lox system (A). Gel electrophoresis of Biopsy (B) and primary microglia DNA (C) from *Tet2*^flox/flox^ and *Tet2*^flox/flox^*Cx3cr1*^Cre/WT^ mice (B). Quantification of Tet2 expression using primers that recognize exon 3 in primary microglia from *Tet2*^flox/flox^*Cx3cr1*^Cre/WT^ and *Tet2*^flox/flox^*Cx3cr1*^WT/WT^ mice with and without 24h LPS treatment (100 ng/ml) (C). Example of line charts for ECAR and OCR generated with Seahorse technology in siRNA control and siRNA Tet2 BV2 microglia cells treated with LPS for 3h (D and E). NOS-2 expression in BV2 cells treated with different doses of 2-deoxy-D-glucose and LPS for 6 h (F and G). Data represented in C and G are shown as mean ± s.d of four independent experiments. Statistics were performed using one-way ANOVA with LSD correction in G. **P* < 0.05, ***P* < 0.01

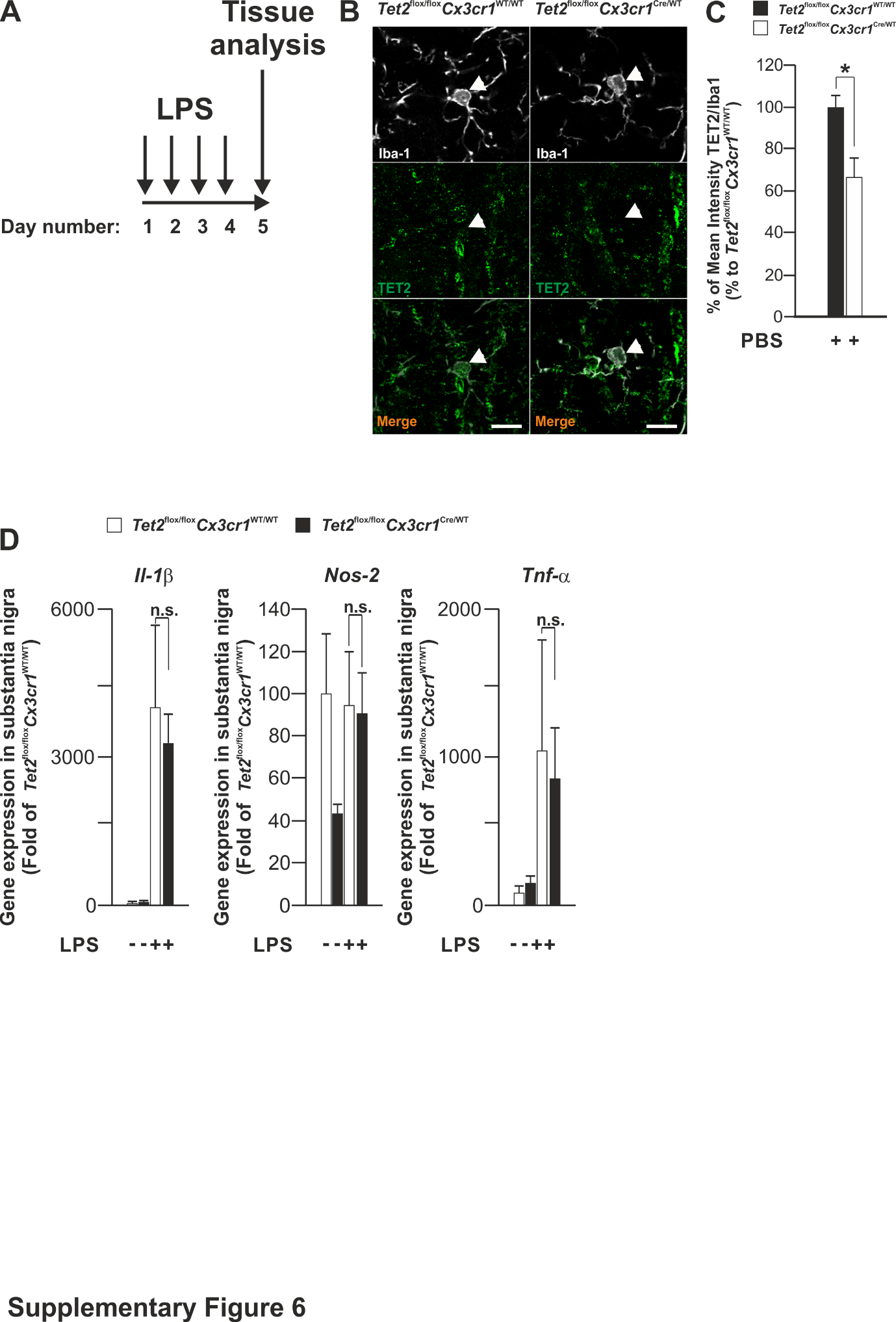

**Figure S6 (related to Figure 6): Effect of intraperitoneal LPS in *Tet2*^flox/flox^ and *Tet2*^flox/flox^*Cx3cr1*^Cre/WT^ mice**

A schematic diagram depicting the number and time of intraperitoneal LPS injections and the day that the mice were sacrificed and their brain tissue analyzed (A). Analysis in vivo of TET2 expression in microglia cells treated only with PBS intraperitoneal injections (following the same pattern of injections as with LPS intraperitoneal injections) (B and C). RT-qPCR for *Il-1β, Nos-2, Tnf-α* expression in substantia nigra of *Tet2*^flox/flox^*Cx3cr1*^Cre/WT^ and *Tet2*^flox/flox^*Cx3cr1*^WT/WT^ treated with LPS or PBS. Data shown as mean ± s.e.m in B and D. The number or independent experiments is three in B and in D, four in *Tet2*^flox/flox^*Cx3cr1*^Cre/WT^ and *Tet2*^flox/flox^*Cx3cr1*^WT/WT^ PBS treated, three Tet2^flox/flox^Cx3cr1^WT/WT^ LPS treated and five *Tet2*^flox/flox^*Cx3cr1*^Cre/WT^ LPS treated, independent experiments. Scale bar in B is 10 μm.

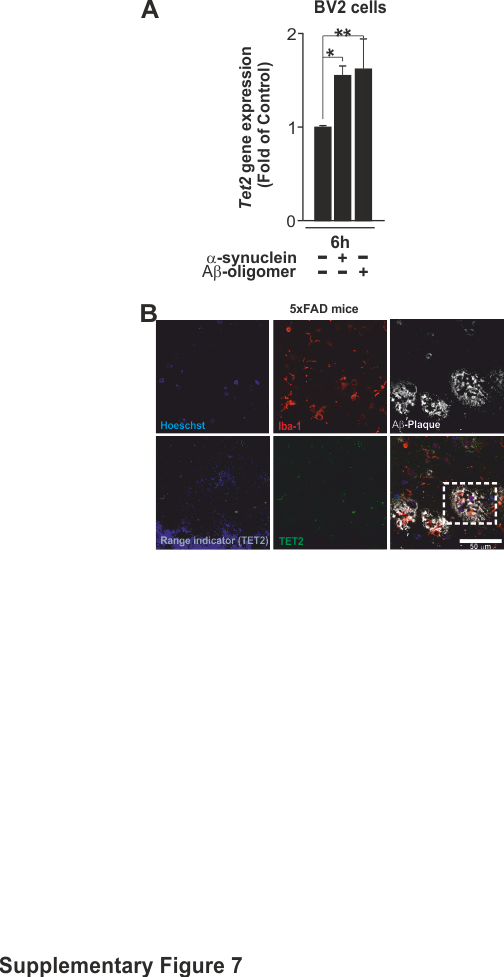

**Figure S7 (related to Figure 7): Microglial TET2 expression upon α-synuclein and Aβ-oligomer treatment and in 5xFAD mice and AD patients.**

*Tet2* expression in BV2 microglial cells after 6 h treatment with α-synuclein fibrils (5µM) and Aβ-oligomers (2 µM). (A). Low magnification of colocalization pictures from Figure 7A (dotted rectangle) in hippocampus of Iba-1, TET2 and β-amyloid in 18 month-old 5xFAD brain sections (B). Data represented as mean ± s.d of three independent experiments for α-synuclein fibrils and four independent experiments for Aβ-oligomers. Statistical analysis was performed using one-way ANOVA with Scheffe correction **P* < 0.05, **P* < 0.01. Scale bar in B is 50 μm
